# Cone Synaptic function is modulated by the leucine rich repeat (LRR) adhesion molecule LRFN2

**DOI:** 10.1101/2023.05.24.542135

**Authors:** Nazarul Hasan, Ronald G. Gregg

**Author notes:** **Author contributions:** NH and RG Designed Research; NH Performed Research and analyzed data; NH wrote and RG edited. **Correspondence to:** Ronald G. Gregg, PhD Department of Biochemistry & Molecular Genetics University of Louisville 319 Abraham Flexner Way Louisville KY 40202. **Conflict of Interest:** The authors declare no competing financial interests. **Funding Sources.** This work was supported by the National Eye Institute (NEI) Grant # EY12354 to RG and NH, and The Preston Joyes Pope Chair in Biochemical Research (RG).

## Abstract

Daylight vision is mediated by cone photoreceptors in vertebrates, which synapse with bipolar cells (BCs) and horizontal (HCs) cells. This cone synapse is functionally and anatomically complex, connecting to 8 types of depolarizing (DBC) BCs and 5 types of hyperbolizing BCs (HBCs). The dendrites of DBCs and HCs cells make invaginating ribbon synapses with the cone axon terminal, while HBCs form flat synapses with the cone pedicles. The molecular architecture that underpins this organization is relatively poorly understood. To identify new proteins involved in synapse formation and function we used an unbiased proteomic approach and identified LRFN2 (Leucine-rich repeat and fibronectin III domain-containing 2) as a component of the DBC signaling complex. LRFN2 interacts with TRPM1 and is selectively expressed at cone terminals and co-localizes with PNA, and other DBC signalplex members. In the absence of LRFN2 the cone-mediated photopic electroretinogram b-wave amplitude is reduced. In LRFN2 deficient mice, the synaptic markers: LRIT3, ELFN2, mGluR6, TRPM1 and GPR179 are properly localized. Similarly, LRFN2 expression and localization is not dependent on these synaptic proteins. These data demonstrate that LRFN2 likely interacts with TRPM1 and its absence compromises normal synaptic transmission between cones and cone DBCs cells.

**Significance Statement:** Signaling between cone photoreceptors and the downstream bipolar cells is critical to normal vision. Cones synapse with 13 different types of bipolar cells forming an invaginating ribbon synapses with 8 types, and flat synapse with 5 types, to form one of the most complex synapses in the brain. In this report a new protein, LRFN2 (Leucine-rich repeat and fibronectin III domain-containing 2), was identified that is expressed on the cones synapses. Using *Lrfn2* knockout mice we show LRFN2 is required for the normal cone signaling.

## INTRODUCTION

Vision begins with the absorption of photons by opsins in rod and cone photoreceptor (PR) cells, which convert photons into a glutamatergic signal at the PR axon terminal. This signal is relayed to post-synaptic bipolar and horizontal cells at the first synapse. The majority of the photoreceptors are rods, which function under dim light conditions. Rod-mediated vision has high sensitivity, but relatively low acuity in humans. Rods make synapses only with depolarizing bipolar cells (DBCs) and horizontal cells (HCs). Cones function under bright light conditions, have lower sensitivity than rods, and in humans mediate our high acuity vision.

Cones connect to 8 types of DBCs and 5 types of hyperpolarizing bipolar cells (HBCs)(West and Cepko, 2022). Cone DBCs, along with HCs dendrites, make invaginating synapses with cone axon terminals and HBCs make flat contacts with cone pedicle base. The bipolar cells make connections with about forty types of retinal ganglion cells (RGC)(Bae et al., 2018), which transmit the signal to the rest of the brain. DBCs use the metabotropic glutamate receptor type 6 (mGluR6) (Nakajima et al., 1993)to detect glutamate concentration in the synapse and modulate membrane potential via the TRPM1 channel(Bellone et al., 2008; Morgans et al., 2009; Shen et al., 2009; Koike et al., 2010). In contrast, HBCs signal through AMPA/kainate-type iGluR receptors (Borghuis et al., 2014; Ichinose and Hellmer, 2016). BCs also can be distinguished based on cellular morphology(Kim et al., 2014; Greene et al., 2016) and gene expression pattern (Shekhar et al., 2016).

Post-synaptic signalplex composition of rod BCs and cone DBC is similar, although their interaction with key organizing molecules such as LRIT3 and ELFNs is different. The loss of LRIT3, results in the loss of nyctalopin and TRPM1 from the signalplex. In contrast, the loss of LRIT3 causes the loss of all the key signalplex proteins, including mGluR6, GPR179, nyctalopin, TRPM1, RGS7/11 and R9AP in cone DBCs. This notable difference between the impact of LRIT3 on rod BC versus cone DBC signalplexes led to the hypothesis that additional unknown synaptic protein(s) are likely required for rod and/or cone DBC signalplex assembly.

In this study, we used a proteomic approach to identify additional synaptic cell adhesion molecules. We identified LRFN2 (Leucine-rich repeat and fibronectin III domain-containing 2) as a TRPM1-interacting protein in mouse retina. LRFN2, also known as SALM1 (synaptic adhesion-like molecule-1), is a type I transmembrane glycoprotein with the N-terminal region that contains leucine rich repeats (LRR), immunoglobulin-like (Ig), and fibronectin type III (Fn3) domains, located in the extracellular space, and the intracellular C-terminal contains a PDZ binding domain (Morimura et al., 2006). Here, we show that LRFN2 is expressed in and localized to the cone terminals, where it colocalizes with the cone terminal marker, PNA. Knockout of LRFN2 does not alter expression of the other signalplex proteins, mGluR6, GPR179, TRPM1 and LRIT3; and conversely they are not needed for normal expression of LRFN2 The lack of LRFN2 has no impact on the scotopic electroretinogram; but does result in a decrease in the photopic b-wave amplitude at the highest flash intensities. This adds another member of a growing group of trans-synapatic proteins that organize the post-synaptic signaling complex in DBCs.

## MATERIALS AND METHODS

### Animals

All procedures were performed in accordance with local Institutional Animal Care and Use Committees and the Society for Neuroscience policies for the use of animals in research. All mice were multiply housed in a local AAALAC-approved facility (Association for Assessment and Accreditation of Laboratory Animal Care) under a 12 hr light/dark cycle. All mouse lines used have been described previously: *Lrit3^-/-^*(Hasan et al., 2019)*, Trmp1^-/-^* (Shen et al., 2009), *Grm6^-/-^* (Masu et al., 1995), and *GPR179^-/-^* (Peachey et al., 2012). These lines were either generated on a C57BL/6J background or backcrossed onto this background for at least ten generations. Mice between the ages of P45 and P100 of either sex were used in the experiments.

### Generation of *Lrfn2^-/-^* and *Elfn2^-/-^* mice with zinc finger nucleases

To create a null allele of *Lrfn2* we contracted with Sigma to develop a ZFN (zibc fngure nuclease) targeted to a highly conserved region in exon 2 of *Lrfn2*. mRNA (10 ng/µl) encoding the ZFNs were injected into 350 C3H/HeNTac/C57BL/6NTac hybrid embryos. 250 viable embryos were implanted into eight Swiss Webster recipient mothers, yielding many offspring. Tail biopsies from offspring were collected, and genomic DNA isolated by digesting tissue in Direct PCR solution (Thermo Scientific) supplemented with 2.5 μg/μl proteinase K (Thermo Scientific). Primers flanking the ZFN target site (Fwd:5’-TAACCTGGGCATAGCCTGTC-3’; Rev: 5’-AAGGTCCAGGAAGGAGAAGG-3’) were used to genotype the mice (WT =439bP, Mutant=401bp). PCR products were sequenced on a 3130xl Genetic Analyzer (Applied Biosystems). The *Lrfn2^-/-^* allele was backcrossed onto C57Bl/6J mice for 10 generations. The mutant allele has a 38bp deletion in exon 2 of the *Lrfn2* gene (chr17:49,070,008-49,070,045 GRCm38/mm10) causing a frameshift mutation that was predicted to be a null allele.

The *Elfn2^-/-^*mouse was created by JAX labs using CRISP/Cas9 technology and has a 7493 bp deletion (chr15:78667284-78674776; GRCm38/mm10) that removes the entire open reading frame for *Elfn2*, which is encoded by a single exon. To genotype the line PCR as described above using primers Elfn2-F: 5’-AGACAGTCCCTACCCACACG-3’; ELFN2-WTR:5’-AGGCTCAGACCTTCAAGCAG-3’; ELFN2-KOR:5’-ACCAGGTTGTCAGCACATCA-3’, yields a 321 bp fragment for the WT allele, and 301bp fragment for the deleted allele.

### Antibodies

Antibodies used in this study are listed in Table 1. The goat anti-mGluR6 antibody was generated commercially (Life Technology Corporation) by immunizing animals with the peptide KKTSTMAAPPKSENSEDAK conjugated to KLH, and affinity purified using the peptide conjugated to a solid support. The specificity of all antibodies were validated on retinal sections from the respective knockout mouse lines.

**Table 1.** Immunohistochemical reagents used in experiments

| Antigen | Dilution | Source | Catalogue # / RRID# |
| --- | --- | --- | --- |
| LRFN2 | 1:1000 | Sigma | SAB3500015; RRID: |
| LRIT3 | 1:1000 | Hasan et al 2019 | N/A |
| GPR179 | 1:2000 | Peachey et al., 2012 | N/A |
| TRPM1 | 1:1000 | Hasan et al 2019 | N/A |
| mGluR6 | 1:1000 | Current paper | N/A |
| Pikachurin | 1:2000 | Wako Chemicals | 011-22631; RRID: |
| PNA | 1:1000 | Molecular probes | L-32460; RRID |
| ELFN2 | 1:2000 | Invitrogen | PA5-43521, RRID |

### Retinal preparation and immunohistochemistry

Mice were euthanized, their eyes were enucleated, and the lens removed. Eyecups were fixed for 15-30 min in 4% paraformaldehyde in phosphate buffer (0.1M PB), pH 7.4. After fixing, the eyecups were washed three times in PBS then cryoprotected in increasing concentrations of sucrose in PB (5%, 10%, 15% for 1 hr each and 20% overnight). Eyecups, usually from multiple genotypes, were embedded in 2:1 OCT/20% sucrose solution and frozen in a liquid nitrogen-cooled bath of isopentane. Eyecups were sectioned (18 µm) using a Leica 1850 cryostat, mounted on Superfrost Plus glass slides (Thermo Fisher Scientific), and stored at −80°C. Before immunostaining, sections were warmed to 37°C and washed with PBS for 5 min and PBS containing 0.05% Triton X-100 (PBX) for 5 min. After blocking in PBX containing 5% normal donkey serum (blocking solution) for 1 hr, sections were incubated with primary antibody diluted in blocking solution overnight at room temperature, and then washed three times for 10 min each with PBX. Sections were incubated with secondary antibody (1:1000) in PBX for 1 hr at room temperature followed by washing for 10 min twice in PBX and once in PBS. Coverslips were mounted to slides using Vectashield (Vector Laboratories). Images were taken using an FV3000 confocal microscope (Olympus) and corrected for contrast and brightness using Fluoview Software (Olympus) or Photoshop (Adobe Systems). Representative images are shown in the figures (typically four sections, each containing multiple genotypes, were placed on a single slide. Retinas from at least three mice were processed and examined. For all experiments the images shown represent data obtained in at least 5 eyes.

### Cell culture, transfection and immunoblotting

Human Embryonic Kidney (HEK293T) cells were cultured in high-glucose DMEM supplemented with 10% fetal bovine serum, 2 mM L-glutamine, 50 IU/ml penicillin, and 50 µg/ml streptomycin. Cells were seeded on 60 mm culture dishes one day prior to transfection and transfected with LRFN2 and TRPM1 expression plasmids using jetPrime reagent (Polyplus transfection) or Lipofectamine 2000 (Invitrogen) according to the manufacturer’s instructions. After 24–48 hr of transfection, cells were harvested in NP-40 lysis buffer (50mM Tris, 150mM NaCl, 2mM EDTA, and 1% Nonidet P40, pH 8.0, supplemented with protease inhibitor cocktail; Sigma-Aldrich).

After rotating for 45 min at 4°C cell debris was removed by centrifugation at 17,000 X *g* for 15 min at 4°C. The supernatant was collected and protein quantified by Bradford reagent (Bio-Rad). Protein lysates were analyzed on 4–12% NuPAGE gels (Invitrogen), transferred to PVDF membranes, and blocked with Intercept (TBS) Odyssey (LI-COR Biosciences). Membranes were incubated with primary antibodies diluted in Odyssey Blocking Buffer and washed four times with TBS containing 0.1% tween-20 (TBST). After incubating with IRDye800 CW and IRDye680 CW-conjugated secondary antibodies diluted in Odyssey Blocking Buffer, membranes were washed four times with TBST. Proteins bands were visualized by scanning the membranes in an Odyssey Infrared Imaging System (LI-COR Biosciences) using both 700 and 800 nm channels.

### Coimmunoprecipitation

For coimmunoprecipitation from HEK293T cells, transfected cell lysates (500 µg total protein in 200 µl) were pre-cleared by incubating with 10 µl Dynabeads protein G (Invitrogen) at 4°C for 1 hr. Pre-cleared lysates were incubated with 2-5 µg of anti-LRFN2 or anti-TRPM1 antibodies overnight at 4°C on an orbital rocker. 30 µl of Dynabeads protein G was added and incubated for 1-2 hr at 4°C. Dynabeads were collected and washed four times with Tris-buffered saline containing 0.3% Tween 20. Protein complexes were eluted with 40 µl of 4X LDS loading buffer by incubation at 70°C for 10 min, separated by SDS-PAGE, and analyzed by immunoblotting.

### Mass spectrometry

Mice were euthanized by CO2 exposure, retinas were isolated from control mice and homogenized in lysis buffer (1% Nonidet P-40, 2 mM EDTA (ethylenediaminetetraacetic acid), and 20 mM HEPES (4-(2-hydroxyethyl)-1-piperazineethanesulfonic acid), pH 7.4, supplemented with protease inhibitor cocktail (Sigma)) by rotating for 45 min at 4°C. Samples were centrifuged at 17,000 × *g* for 20 min at 4°C to remove the cell debris, and supernatant was pre-cleared with Dynabeads (Invitrogen) for 1 hr at 4°C. Samples were incubated with anti-LRFN2 or anti-TRPM1 antibodies overnight at 4°C. Dynabeads were added to lysates and incubated for 1.5 hr at 4°C. Protein complexes were eluted from Dynabeads with NuPAGE LDS sample buffer (Invitrogen) and electrophoresed on NuPAGE gels (Invitrogen), until the highest molecular weight standard (260 kDa) had moved ∼5 mm into the gel. Electrophoresed gel pieces were cut from the top of the gel and an in-gel tryptic digestion was performed.

The resulting peptide mixture was resolved by liquid chromatography (LC) using an EASY n-LC (Thermo Scientific) UHPLC system with buffer A (2% v/v acetonitrile/0.1% v/v formic acid) and buffer B (80% v/v acetonitrile/0.1% v/v formic acid) as mobile phases. The mass spectrometry data from LC elutes was collected using an Orbitrap Elite ETD mass spectrometer (Thermo Scientific). A decision tree was used to determine whether CID or ETD activation was used.

Proteome Discoverer v1.3.0.330 was used to analyze the data collected by the mass spectrometer. Scaffold software (version 4.10.0) was used to calculate the false discovery rate using the peptide and protein prophet algorithms.

### Electroretinography (ERG)

Mice were dark-adapted overnight and anesthetized with ketamine (117.5 mg/kg) and xylazine (11.3 mg/kg) solution prepared in Ringer’s solution, and prepared for ERG recordings using dim red light. Pupils were dilated with topical applications of 0.625% phenylephrine hydrochloride and 0.25% Tropicamide and the corneal surface anesthetized using 1% proparacaine HCL. Body temperature was maintained via an electric heating pad (TC1000 Temperature control, CWE Inc.). ERGs were recorded using a clear contact lens with a gold electrode contacting the corneal surface wetted with 1% methylcellulose. Needle electrodes in the tail and on the midline of the forehead serve as a ground and reference, respectively. Scotopic responses were recorded by presenting light flashes (from −3.6 to 1.4 log cd sec/m^2^) to dark adapted mice. For photopic ERGs, mice were light adapted to 20 cd/m^2^ for 5 min and responses were measured by presenting light flashes (from −0.8 to 1.4 log cd sec/m^2^) on this rod saturating background.

### Statistical analyses

Prism 9.5.0 software (Graphpad Software, Inc., La Jolla, CA) was used to perform the appropriate statistical analyses (see text and figure legends) for the necessary comparison. Statistical significance was determine at p≤ 0.05.

## RESULTS

TRPM1 is a non-specific cation channel that is part of a signalplex that includes mGluR6 and GPR179 and is critical to the function of DBCs. To identify additional TRPM1 interacting partners we designed a proteomic screen, using TRPM1-specific antibodies to immunoprecipitate the TRPM1 complex from protein lysates prepared from control mouse retinas. As a control we used a non-immune IgG to immunoprecipitate proteins from control retina lysates. Mass spectrometry of the immunoprecipitates resulted in the identification of thousands of peptides that could be mapped to hundreds of proteins. Analysis of proteins immunoprecipitated specifically with the TRPM1 antibody revealed the identification of a known interacting partner, GPR179 (Ray et al., 2014), in addition to TRPM1 itself (48 unique peptides; 37% coverage) (Fig 1A). We then filtered the results for cell adhesion molecules and identified LRFN2 (leucine rich repeat and fibronectin type III domain containing 2), which was identified by seven unique peptides (coverage of 13% of the amino acid sequence (Fig 1A,C). To further evaluate whether LRFN2 was part of the TRPM1 complex we performed immunoprecipitations with a LRFN2 antibody followed by proteomic analyses. These analysis identified 21 unique LRFN2 peptides (36% coverage) and 4 unique peptides that matched TRPM1 (3.4% sequence coverage) (Fig 1B). No TRPM1 or LRFN2 peptides were identified from IgG controls. Based on these data we hypothesize that TRPM1 and LRFN2 are part of the same signaling complex and likely interact directly.

**Figure 1.**
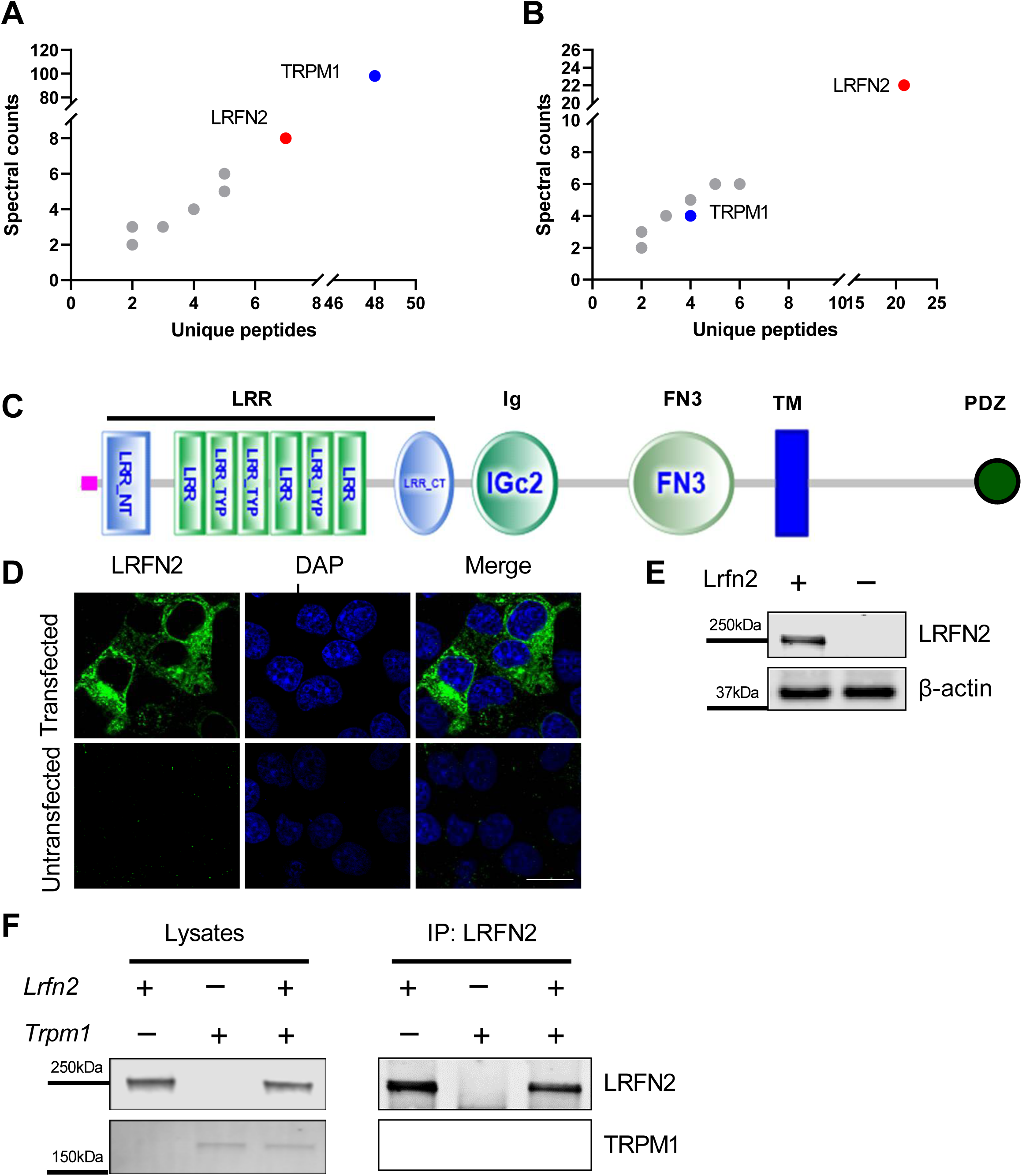
LRFN2 is an interacting partner of TRPM1 in mouse retina. Analysis of proteins co-purified with TRPM1 (A), LRFN2 (B) and identified by mass spectrometry in wild type mouse retina. (C) Domain composition of LRFN2 predicted by SMART® program. LRFN2 contains a signal sequence (purple box), extracellular LRR, Ig and FN3 domains, a transmembrane segment (TM, blue box), and intracellular C terminus containing a PDZ binding domain (blue ball). (C,D). Immunohistochemistry (C) and western blots (D) showing expression of LRFN2 expressed in HEK293T. (F) LRFN2 and TRPM1 interact *in vitro.* Western blots of lysates (left panel) or immunoprecipitates (right panel), after expression of combinations of LRFN2 and TRPM1 in HEK293T cells. Proteins were co-immunoprecipitated with LRFN2 antibodies.

### LRFN2 interacts with TRPM1 in the HEK293T cells

LRFN2 contains leucine-rich repeats, immunoglobulin and fibronectin domains (Fig 1C), which are known to be involved in protein:protein interactions. To determine if LRFN2 and TRPM1 interact directly we expressed one or both in HEK293T cells and did co-IP experiments.

HEK293T cells were transfected with PGK-LRFN2 and/or CMV-TRPM1 plasmids and protein expression confirmed was by immunocytochemistry (Fig 1D) and western blotting (Fig 1E,F). To determine if LFRN2 and TRPM1 interacted, we expressed one or both and performed co-IP using antibodies against LRFN2. Western blot analyses of single-and double-transfected cell lysates show that both proteins were expressed in lysates (Fig 1F). In cells only expressing LRFN2 the IP showed presence of LRFN2 only. When both LRFN2 and TRPM1 were expressed, co-IP with LRFN2 antibodies showed the presence of TRPM1 (Fig 1F). These results combined with the mass spectrometry data indicate that LRFN2 and TRPM1 are part of the same protein complex and likely interact directly.

### LRFN2 is expressed on cone pedicles in mouse retina

To determine the location and function of LRFN2 in retina, we generated a *Lrfn2^-/-^* mouse using zinc finger nucleases. Several lines containing indels were created and we characterized one that had a 38 base pair deletion within exon 2. This created a frameshift mutation and was expected to be a null allele. To determine the expression pattern of LRFN2 in retina we used a commercially available antibody (Sigma, Cat# SAB3500015). To verify its specificity we compared staining of control and *Lrfn2*^−^*^/^*^−^ retinas, using both immunohistochemistry and western blotting (Fig 2A-B). The LRFN2 antibody showed staining exclusively in the OPL of the control retina, and was absent in the *Lrfn2*^−^*^/^*^−^ retina (Fig 2A,C). Western blots also showed that the antibody was specific to LRFN2, staining a 250 kd band in control that was absent in the *Lrfn2^-/-^*retina lysates (Fig 2B).

**Figure 2.**
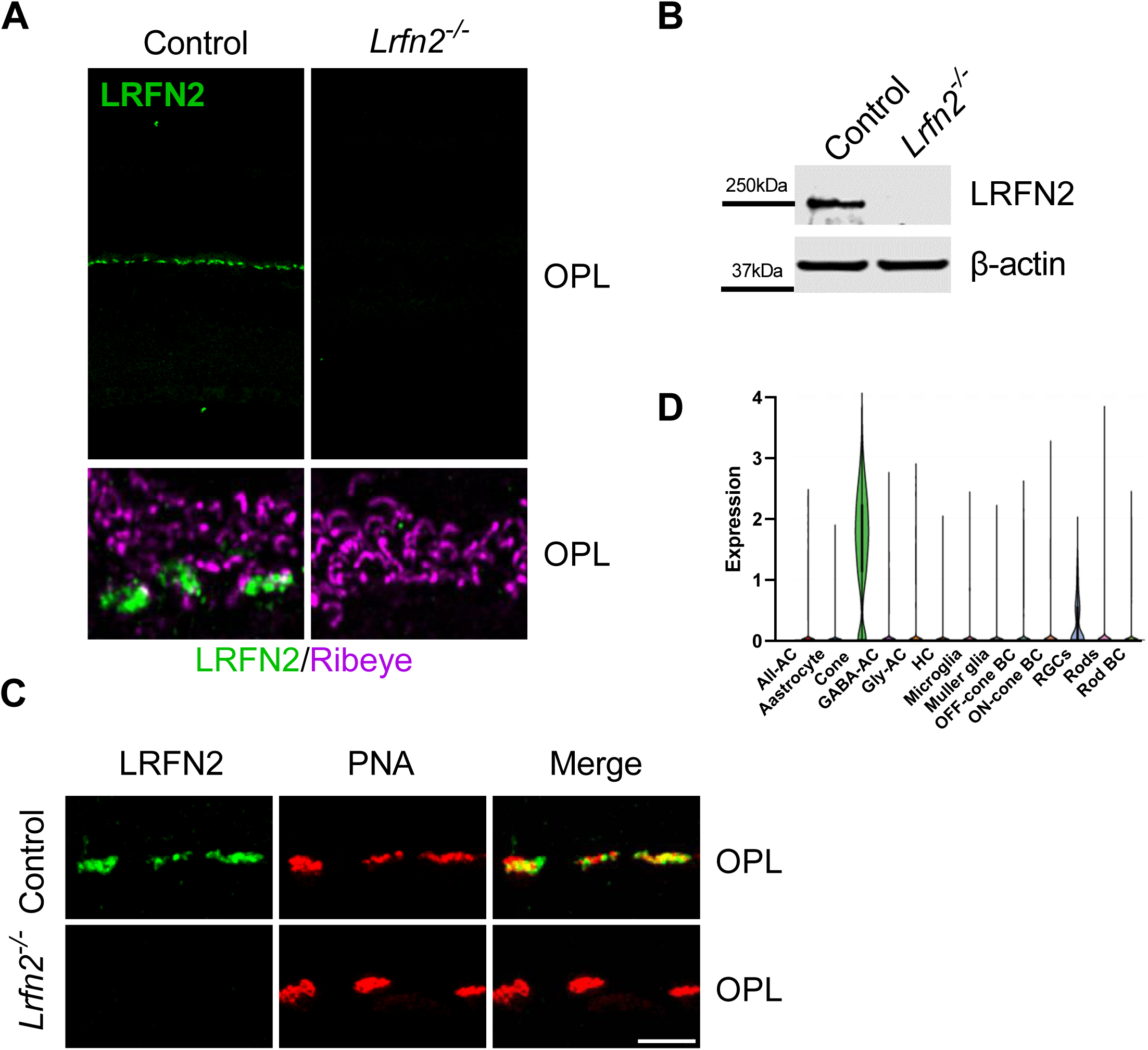
LRFN2 is localized to cones synapses in the OPL of mouse retina. (A) Representative confocal images after immunohistochemical staining for LRFN2 (green) and Ribeye (magenta) in retina slices from control (left panels) and in *Lrfn2^-/-^* retinas (right panels). Scale bar = 20µm. (B) Western blot showing absence of LRFN2 in *Lrfn2^-/-^* retinas. β-actin as loading control. (C) Immunohistochemical localization LRFN2 (green) with the cone terminal maker PNA (red) in retinal sections from Control and *Lrfn2^-/-^* retinas. (D) scRNAseq of retina showing expression of LRFN2 in cones, and less abundantly in RGCs (data from (Wang et al., 2022)).

The staining pattern in the control retina OPL resembles that seen for the cone terminal marker PNA (Blanks and Johnson, 1983). To evaluate this we double labelled transverse retinal sections for LRFN2 and for PNA. Figure 2C shows that LRFN2 and PNA co-localize. Because of the imaging resolution it is not possible to determine if LRFN2 is expressed in cones or cone DBCs. To address this question we used publically available single cell RNA sequencing data from retina (Wang et al., 2022) (Fig 2D). These data show that LRFN2 expression is enriched in cones with some expression in RGCs. Thus, because it interacts with TRPM1 this indicates that LRFN2 likely acts in a trans-synaptic manner, similar to LRIT3 and ELFN2 (Hasan et al., 2019; Cao et al., 2020; Hasan et al., 2020). Combined, these results support the conclusion that LRFN2 expression is specific to cones and the protein is localized to cone terminals.

### ON bipolar cell signalplex protein expression in *Lrfn2*^−^*^/^*^−^ retina

Previous studies analyzing various knockout mouse lines have shown that there is a complex relationship and interdependency between the expression of DBC signalplex proteins(Gregg et al., 2014). For example, LRIT3 is required for localization of mGluR6, TRPM1, nyctalopin, GPR179, and RGS7/11 in cone DBCS (Neuille et al., 2015; Neuille et al., 2017; Hasan et al., 2020). Similarly, nyctalopin is required for TRPM1 insertion at the DBC dendritic tips (Pearring et al., 2011). These data show that signalplex proteins are interdependent for their localization and function. To examine if LRFN2 was required for and dependent on other DBC signalplex proteins we examined expression in knockout mice. We created *Lrfn2^-/-^* mice and used immunohistochemistry to examine expression of TRPM1, LRIT3, mGluR6, and GPR179, Pikachurin, Ribeye and ELFN2, which are expressed at cone to cone DBC synapses. In all cases the expression pattern in the *Lrfn2^-/-^* mice was indistinguishable from controls (Fig. 3,4) To determine if the expression of LRFN2 was dependent on LRIT3, TRPM1, mGluR6, GPR179, Nyctalopin, or ELFN2, we determined its expression pattern in *Grm6*^−^*^/^*^−^*, Trpm1*^−^*^/^*^−^,GPR179*^-/-^*, *Nyx^nob^*, and *Elfn2^-/-^* knockout mouse lines, respectively (Fig. 5). To mark the cone terminals in each section we stained with PNA. In all these knockout lines the staining pattern for LRFN2 in *Lrfn2^-/-^* retinas was indistinguishable from controls. Combined these data showed that the expression of LRFN2 was not dependent on the expression of any post-(mGluR6, TRPM1, GPR179, Nyctalopin) or pre-synaptic (LRIT3 or ELFN2) protein tested. Therefore, even though LRFN2 and TRPM1 interact their localization to the DBC dendritic terminals is not interdependent, nor is there any interdependency of LRFN2 with other DBC signalplex members.

**Figure 3.**
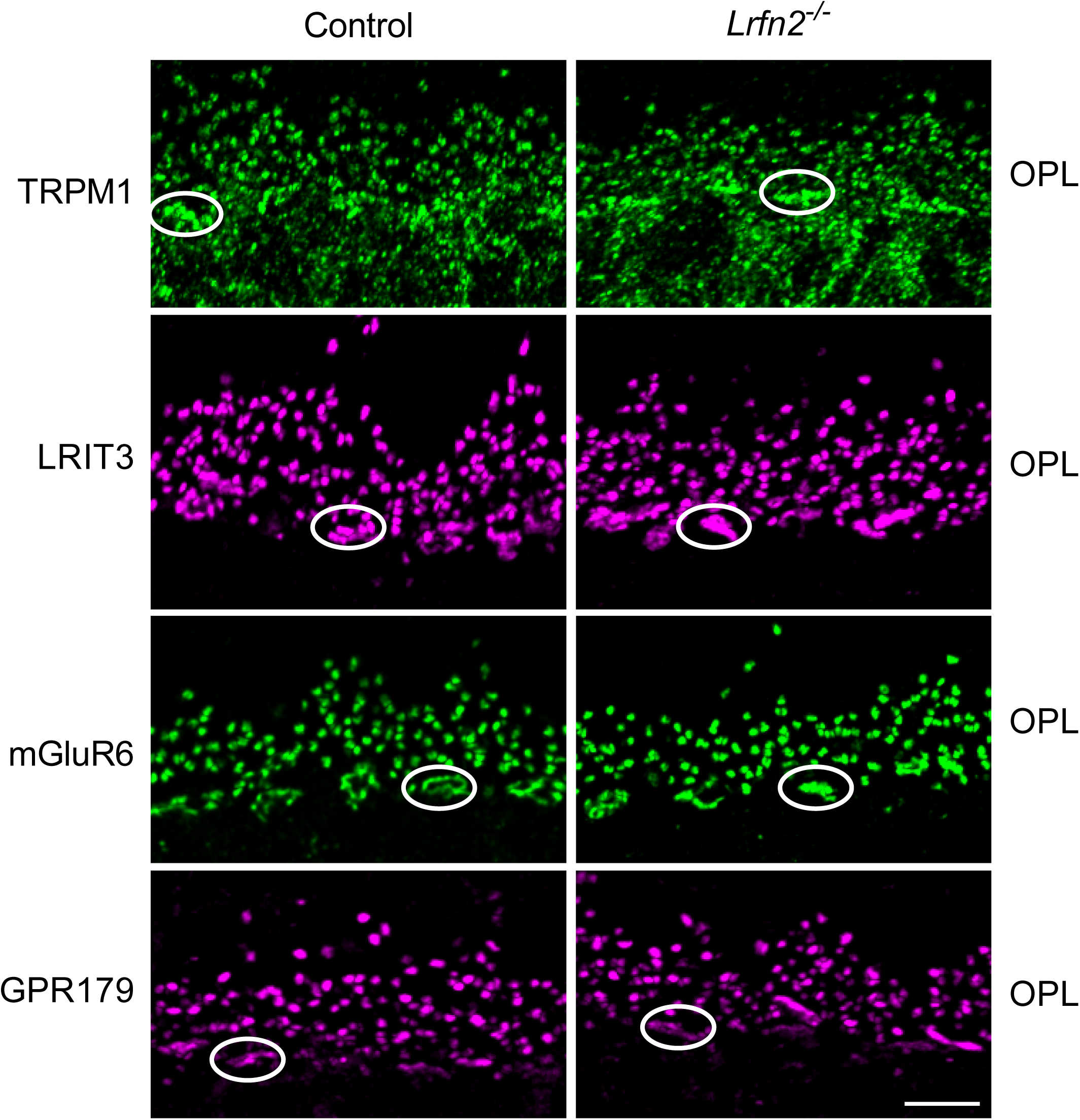
Synaptic proteins are localized normally in *Lrfn2^-/-^* mouse retina. Representative confocal images showing immunohistochemical localization of post-synaptic synaptic proteins TRPM1, LRIT3, mGluR6 and GPR179 in retinal slioces from Control and *Lrfn2^-/-^* retinas. Scale bar = 5µm. Circles indicate cone synapses. OPL, outer plexiform layer.

**Figure 4.**
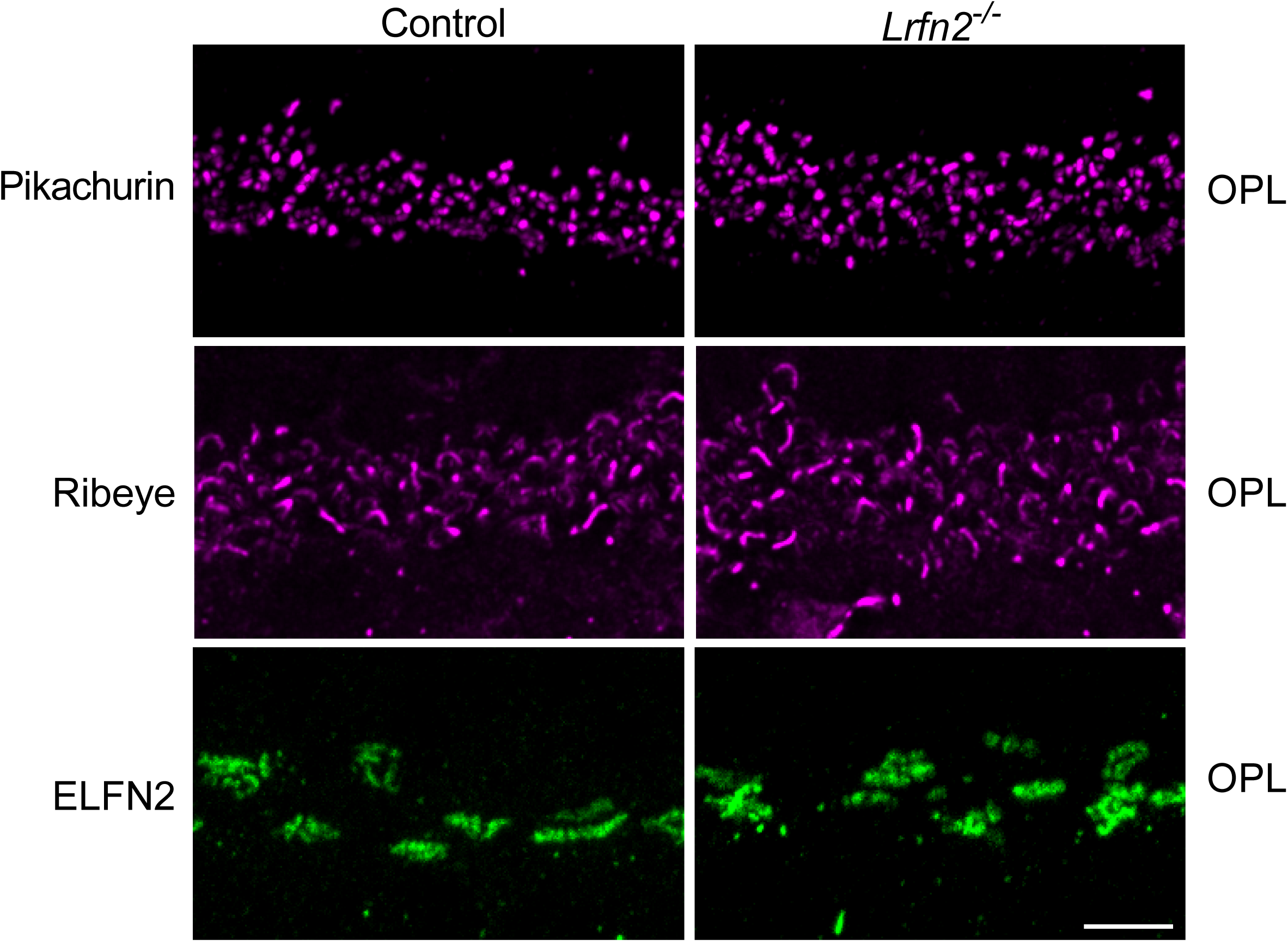
Synaptic proteins pikachurin, ribeye and ELFN2 are localized normally in *Lrfn2^-/-^* mouse retina. Note the ELFN2 is only expressed on cone terminals. Scale bar = 5µm.

**Figure 5.**
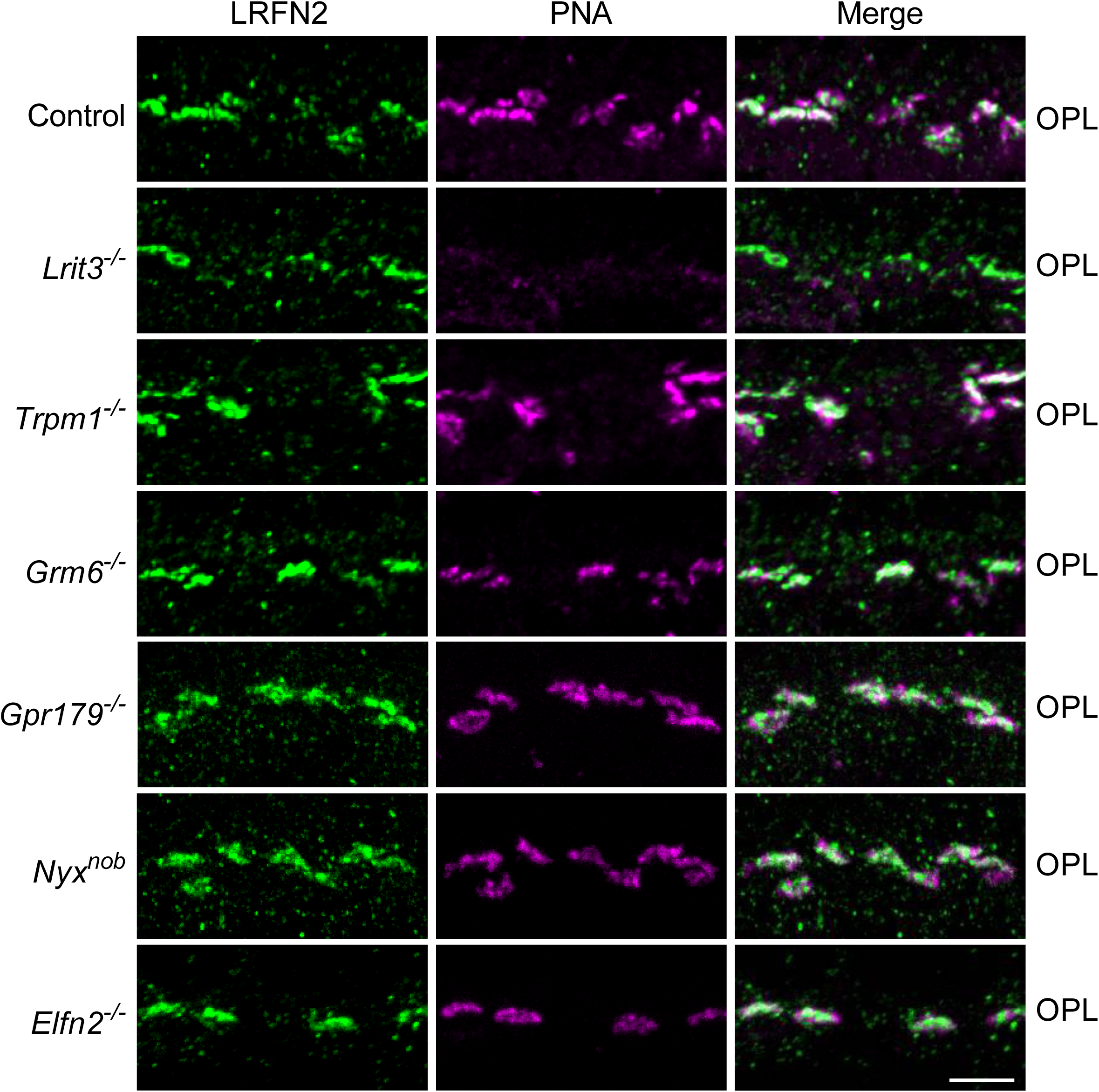
LRFN2 is expressed normally in *Lrit3^-/-^, Trpm1^-/-^*, *Grm6^-/-^*, *Grm6^-/-^*, *Gpr179^-/-^*, *Nyx^nob^* and *Elfn2^-/-^* retinas. Images are from representative retinal sections stained for LRFN2 (green) and PNA (magenta). Scale bar = 5µm. OPL, outer plexiform layer.

### Function of LRFN2 in signal transduction in mouse retina

To assess the possible impact of loss of LRFN2 on function we used the electroretinogram (ERG), which assesses photoreceptor, and connected DBCs function. Recordings were done at 6-8 weeks of age under either scotopic or photopic conditions, to assess rod BC and cone DBC function, respectively. The waveforms for several flash intensities in a control and *Lrfn2^-/-^* mouse are shown under scotopic (Fig 6A) and photopic (Fig 6B) conditions. Summary data for control (n=7) and *Lrfn2^-/-^* (n=8) are shown in Fig 6C and 6D, respectively. As expected given LRFN2 is not expressed at the rod to rod BC synapse, the a-wave or b-wave amplitudes of the electroretinogram under scotopic conditions in *Lrfn2^-/-^* mice were not different than controls (Fig 6A, C). The photopic ERG, which reflects cone-pathway function, showed a normal a-wave at all flash intensities (Fig 6B,D). In contrast, the b-wave amplitudes of the photopic ERG in *Lrfn2*^−^*^/^*^−^ mice were significantly reduced at the 3 highest flash intensities tested Fig. 6D. (2-way ANOVA, adjusted for multiple testing using the Sidak correction). The estimation plots of the 95% CI of the difference between controls and *Lrfn2^-/-^* for a-and b-waves, for scotopic and photopic conditions are shown in Extended Figure 5-1. This reduction in photopic b-wave amplitudes indicates an alteration in signal transduction between cones and cone DBCs.

**Figure 6.**
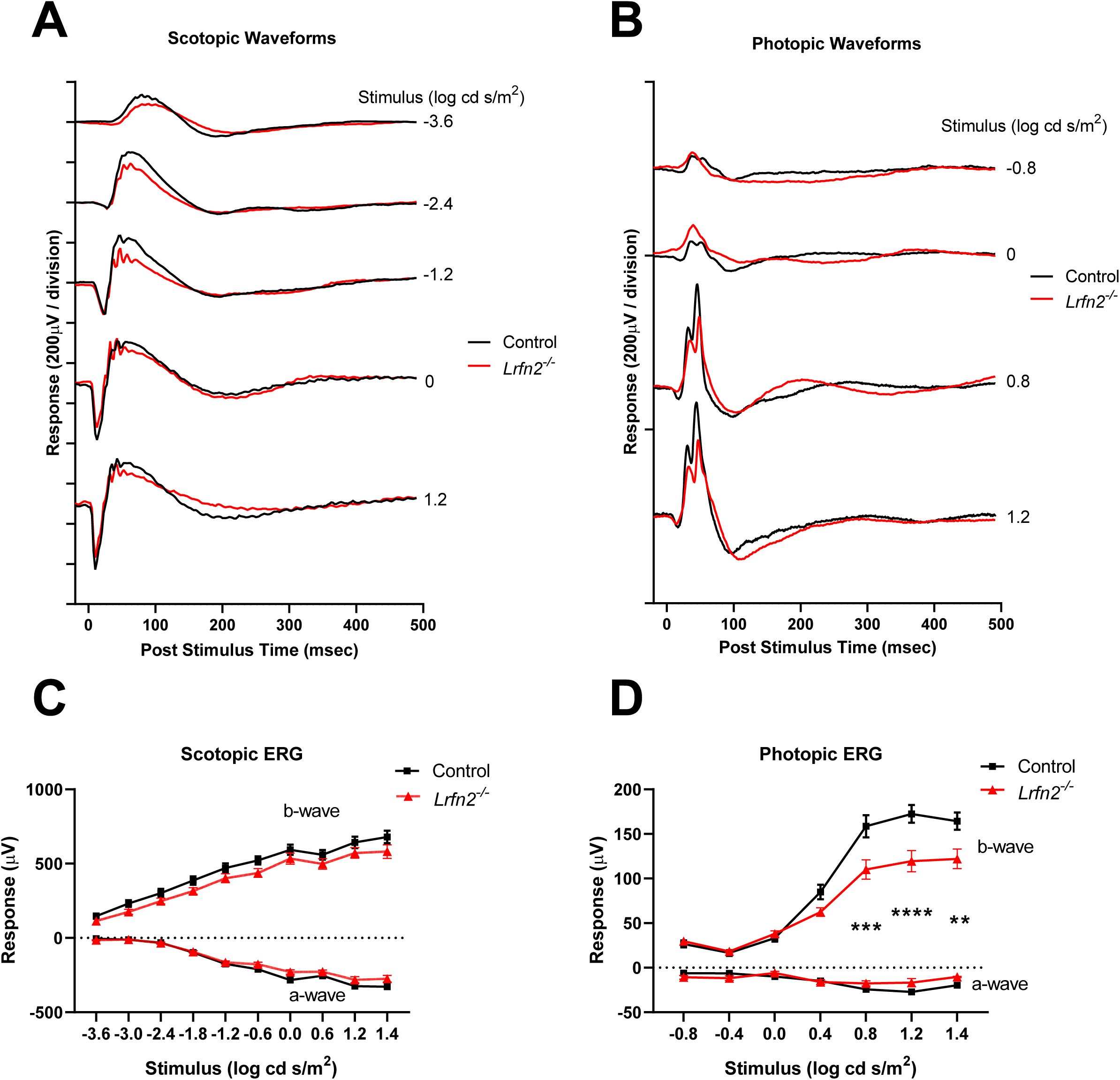
Photopic ERG b-waves of *Lrfn2^-/-^* mice have a reduced b-wave. (A,B) Waveforms for a single animal under scotopic (A) and photopic (B) ERG responses at different intensity flashes for one control (black) and one *Lrfn2^-/-^* eye (red). Average stimulus-response plots for the ERG a-wave and b-wave amplitudes under scotopic (B) and photopic (D) conditions for control (n=7, black) and *Lrfn2^-/-^* (n=8, red). Statistics: (D) comparison of control vs *Lrfn2^-/-^* groups, ** padj<0.01, *** padj<0.001 for stimuli, 2-way repeated measures ANOVA. Between the groups, there are no differences in b-wave amplitude under scotopic conditions, and in a-wave amplitude under either scotopic or photopic conditions (2-way repeated measures ANOVA, padj>0.05). 95% confidence intervals of the differences between genotypes are shown in Extended Figure 1.

## DISCUSSION

Here we show that LRFN2 (Leucine rich repeat and fibronectin type III domain containing 2) is part of the cone DBC signalplex, and represents another member of the trans-synaptic proteins required for normal visual function. LRFN2 contains LRR, Ig and Fn3 domains predicted to be extracellular. The intracellular domain of ∼130 amino acids also contains a PDZ binding domain (Morimura et al., 2006). Cones make synapses with, and relay high-sensitivity signals to many subtypes of ON and OFF cone bipolar cells. However, the molecular components required for synapse formation, neurotransmitter release and its detection by correctly localized post-synaptic receptor complexes is poorly understood. Our proteomic approach identified LRFN2 as another novel DBC signalplex interacting proteins.

LRFN2 has been studied in the brain (Thevenon et al., 2016; Morimura et al., 2017; Li et al., 2018; McMillan et al., 2021), and is a presynaptic organizer of synapse development by promoting F-actin/PIP2-dependent clustering of Neurexin in hippocampal neurons (Brouwer et al., 2019). In *Lrfn2^-/-^* mice there is a decrease in synaptic plasticity and inhibitory synapse development (Li et al., 2018). These mice also have altered behaviors, specifically decreased acoustic vocalization of pups and increased startle response, although there was no effect on locomotion. A microdeletion in humans containing the *LRFN2* gene was found associated with selective working memory deficits, and borderline intellectual functioning (Thevenon et al., 2016).

LRFN2 associates with and regulates surface expression of AMPA receptors, synaptic activity and hippocampal long-term potentiation through interaction with nexin-27 (McMillan et al., 2021). Whether LRFN2 interacts with nexin’s at the cone terminal is unknown. Here we show LRFN2 interacts with TRPM1, but its expression is restricted to cone terminals. The single cell RNA sequencing data (Peng et al., 2019; Wang et al., 2022) suggest its expression is in cones in the outer retina. This is consistent with the fact that LRFN2 contains a PDZ binding domain (Morimura et al., 2006) that could interact with PSD95, which is a presynaptic protein in the retina (Koulen et al., 1998). Thus LRFN2, like LRIT3 and ELFN2 serve as trans-synaptic scaffolds between cones and cone DBCs.

At the functional level the loss of LRFN2 causes no change in the scotopic ERG, consistent with is expression only on cones. However, it is required for the normal signaling of cone DBCs at bright flash intensities. Once question that arises is that given LRFN2’s interaction with TRPM1 why the phenotype is so subtle. One possibility is that because the *Lrfn2^-/-^*mouse lacks LRFN2 during development that there is compensation by another LRFN family member (LRFN1,3,4, or 5). scRNA seq studies show gene expression of all five members in both mouse and human retina (Balasubramanian et al., 2021; Wang et al., 2022), although currently their localization and developmental expression pattern is unknown. Such compensation was recently shown to occur for another cone-specific trans-synaptic protein, ELFN2(Cao et al., 2022), the loss which had no effect on the synaptic function as measured by the ERG. However, this was because ELFN1 was used to compensate for its absence (Cao et al., 2022). Whether a similar type of compensation occurs for loss of LRFN2 remains to be determined.

In conclusion, our studies have identified a new LRR containing protein, LRFN2, that likely acts in a trans-synaptic manner between cone terminals and the cone DBC signaling complex to modulate the function of TRPM1, and thus signaling through cone DBCs LRFN2 is a member of a large family of LRR-containing proteins that includes FLRTs, NGLs, Slitrks, LRRTM and SALMs, which are expressed throughout the CNS and many have been shown to impact retinal synaptic structure and function(Schroeder and de Wit, 2018). The detailed method of action of LRFN2 remains for future studies.

## Acknowledgements

This work was supported by funding from the NIH

