## Supplementary figures and images for "Cone Synaptic function is modulated by the leucine rich repeat (LRR) adhesion molecule LRFN2"

### Figure 5-1

#### Confidence Intervals (95%) of differences between Control and *Lrfn2*$^{-/-}$ mice

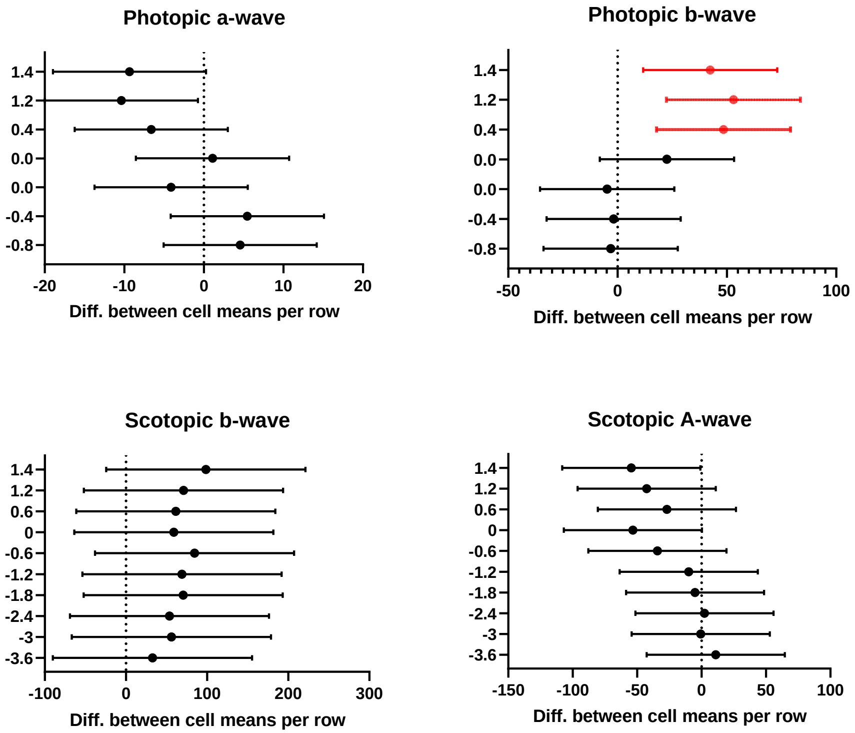
